# Amplitude without phase: interpersonal movement coordination grades inter-brain coupling

**DOI:** 10.64898/2026.09.09.750441

**Authors:** Noor Tasnim, Rachel M. DeLauder, Mackenzie Aychman, Zachary Moore, Mary Gahagan, Jessica Purevtugs, Jesse Newpol, Ryan Frank, Grace Grizzell, Grace Nobriga, Sarah Rose, Julia C. Basso

## Abstract

Interpersonal movement coordination is a foundational feature of human sociality, scaffolding communication, affiliation, and joint action from infancy onward. Whether this behavioral coupling is accompanied by genuine coupling between brains — rather than one manufactured by shared sensory input or motion artifact — has proven difficult to establish. We recorded dual-brain EEG with synchronized head-mounted accelerometry from participant–partner dyads (16 dyads, 155 dyad-condition recordings) across six conditions grading interpersonal coordination from stillness to fully reciprocal joint movement, using improvisational dance to elicit sustained, naturalistic, gradable coordination. Windowed cross-correlation confirmed graded, partner-specific movement coordination, including a lead–lag signature that reversed with who led. Against a pseudo-dyad surrogate, inter-brain synchrony took one specific form: amplitude-envelope coupling in the high-beta and gamma bands, robust to a leakage-corrected measure, with no phase-based or slow-band coupling. Within dyads, movement synchrony predicted the strength of this amplitude coupling (standardized β = 0.54 in gamma), an association that strengthened monotonically with frequency, held under a dyad fixed-effects estimator, and was undiminished when the amount of movement was entered into the model. Movement synchrony and inter-brain coupling also co-fluctuated within interactions, above surrogate. Two control analyses excluded the leading alternatives: the coupling was not sensorimotor rhythm-tracking, since corticokinematic coherence was present but non-localized and not movement-selective, and not a shared movement-frequency rhythm linking the brains. These results identify interpersonal movement coordination as an organizing variable for inter-brain coupling and specify that coupling as fast-band amplitude co-modulation, not phase synchrony. Brains couple when bodies coordinate, not when they merely move together.

## 1. INTRODUCTION

Human social life is built on coordination. From the turn-taking in conversation to the synchronized movements in ritual, music, and dance, people continuously align their actions with one another, and this interpersonal coordination is among the most reliable behavioral signatures of social engagement(Schmidt and Richardson, 2007). Moving in synchrony is not merely a byproduct of shared activity: experimental work shows that it promotes affiliation, trust, cooperation, and prosocial behavior(Rennung and Göritz, 2016; Wiltermuth and Heath, 2009), and that spontaneous movement coupling (head nods, gaze alignment, postural matching) is stronger between individuals with closer relationships(Bell, 2020; Bernieri, 1988; Feldman, 2007; Kinreich et al., 2017), ultimately forming the substrate of nonverbal communication(Bernieri and Rosenthal, 1991). Evolutionary accounts propose that this capacity was selected because coordinated movement strengthened social bonds and signaled coalitional strength among individuals acting together(Hagen and Bryant, 2003; Savage et al., 2020). Interpersonal movement coordination is thus a foundational feature of human sociality — one whose behavioral consequences are well documented, but whose neural basis remains only partially understood(Redcay and Schilbach, 2019).

A natural question is whether coordination between bodies is accompanied by coordination between brains. Hyperscanning, the simultaneous recording of neural activity from two or more interacting people, has revealed inter-brain synchrony across many social contexts, including caregiver–infant exchange(Li et al., 2024), joint music-making(Abalde et al., 2024), and shared attention(Koike et al., 2016). Such coupling is often strongest when partners can see and respond to one another(Hirsch et al., 2017; Leong et al., 2017), and studies that measure behavior alongside brain activity have linked spontaneously coordinated movement to the degree of inter-brain synchronization(Dumas et al., 2010). In the most direct demonstration to date, dyads instructed only to look at one another nonetheless synchronized their body movement, smiling, and eye contact, and each of these behaviors predicted the strength of inter-brain coupling(Koul et al., 2023). What remains untested is whether coupling tracks interpersonal coordination when coordination is itself manipulated rather than left to arise spontaneously — whether it scales with the structure of a jointly performed task, and whether it depends on which partner leads. However, a persistent interpretive problem shadows this literature: when two brains appear synchronized, it is difficult to determine whether the coupling reflects genuine social interaction or is instead produced by shared sensory input, common task structure, or correlated movement and its attendant artifacts(Schilbach and Redcay, 2025).

This problem is not incidental; it is the central methodological challenge of the field. Because interacting partners share an environment, hear the same sounds, and often move in similar ways, any observed inter-brain synchrony has multiple candidate sources. Recent perspectives have therefore argued that interpreting inter-brain coupling requires paradigms capable of dissociating genuine interaction from these confounds, ideally by measuring interpersonal behavior directly and testing whether neural coupling tracks it(Schilbach and Redcay, 2025). Movement is central to this challenge in two ways: it is a primary driver of correlated brain activity between partners, and it is simultaneously a leading source of motion and muscle artifacts that can masquerade as neural coupling. Distinguishing coupling that is organized by interpersonal movement coordination from coupling that is merely a correlate of shared movement therefore requires measuring movement and brain activity together, with surrogate controls that isolate what is specific to the actual pairing.

Partnered dance provides an unusually powerful setting in which to address these questions. Unlike conversation or incidental interaction, dance elicits sustained, whole-body, reciprocal coordination that can be graded in structure, from imitative mirroring to open reciprocal exchange, while remaining naturalistic. Improvisational dance, in particular, cannot be planned in advance: it is a form of unplanned coordination, a collective achievement accomplished without a predetermined plan, fixed roles, or a leader, in which each partner remains autonomous even while acting jointly(Hasan and Kayle, n.d.). Because there is no predetermined plan, improvisation demands continuous, real-time attention and adaptation to a partner, placing interpersonal attunement at the center of the activity — a shift from individual action toward a shared “we-agency,” grounded in perceiving and responding to others moving in a common space(Himberg et al., 2018). These features make partnered improvisational dance not a niche subject but an ideal experimental probe of interpersonal coordination: it produces rich, gradable, reciprocal movement of exactly the kind whose neural signature is in question.

Hyperscanning studies of coordinated dyadic action, though still scarce, are suggestive: inter-brain phase coupling rises during rhythmic perturbations in musical duets(Lender et al., 2023), and in the few EEG studies of partnered dance, neural signals track coordination-relevant features of the interaction, including effects of visual contact(Bigand et al., 2025). Advances in mobile EEG have made naturalistic recording feasible, including proof-of-concept capture of dancers during live performance(Theofanopoulou et al., 2024). Neural synchrony has also been observed among dance audiences, where band-limited coupling tracks engagement(Rai et al., 2025); notably, however, this audience coupling reflects shared sensory stimulation rather than reciprocal interaction(Schilbach and Redcay, 2025) — precisely the distinction that makes interactive, movement-resolved paradigms necessary. The whole-body, reciprocal, role-differentiated coordination that partnered dance affords, measured alongside the movement that produces it, remains largely unexplored.

One framework makes an explicit prediction about how coordinated movement should shape brain activity across people. The Synchronicity Hypothesis of Dance proposes that moving together aligns neural activity not only within a single brain, across the systems supporting sensory, motor, cognitive, social, emotional, and rhythmic processes(Basso et al., 2020), but also between brains, such that interpersonal movement coordination gives rise to inter-brain synchrony. This hypothesis reframes inter-brain coupling as a consequence of coordinated action rather than an incidental correlate of it — and, crucially, it is testable: if movement coordination organizes inter-brain coupling, then the degree of coupling should track the degree of interpersonal coordination, moment to moment, in a manner specific to the interacting pair and not explained by movement amount alone.

Prior work has shown that inter-brain coupling tracks behavioral synchrony as it arises spontaneously; whether it tracks interpersonal coordination when coordination is itself manipulated rather than left to vary has not been tested. Here we test that prediction directly. Using dual-brain hyperscanning EEG recorded simultaneously with whole-body accelerometry, we measured interpersonal movement coordination and inter-brain coupling in participant–instructor dyads across six conditions that parametrically graded interpersonal coordination, from a resting baseline through non-movement social conditions (eye gaze, conversation) to reciprocal joint movement (mirroring and improvisation). Improvisational dance served as the means of eliciting sustained, gradable, reciprocal movement; the questions and measures concern interpersonal coordination and its neural signature in general. Throughout, pseudo-dyad surrogates — pairing each participant with mismatched partners performing the same task — isolated coupling specific to the actual pairing from coupling attributable to shared task structure. Simultaneous accelerometry then allowed us to test whether neural coupling tracked interpersonal movement, while controlling for the amount of movement itself(Burgess, 2013). We asked, first, whether interpersonal movement coordination and inter-brain coupling are each present and partner-specific; second, whether movement coordination predicts inter-brain coupling within dyads, and if so, in which neural measures and frequency bands; and third, whether the two co-fluctuate moment-to-moment within interactions, and whether either leads the other. In addressing these questions, we sought to distinguish inter-brain coupling that is genuinely organized by interpersonal coordination from synchrony attributable to shared sensory input or movement-related artifacts.

## 2. METHODS

### 2.1 Participants and study design

Participants were 18 adults (83.3% female; age range 18–45 years) with no or limited prior dance experience, recruited through community outreach in the New River Valley (social media, dance-class listings, and flyers at local businesses). Inclusion criteria were age ≥ 18 years, ability to engage in physical activity, and English proficiency; exclusion criteria were pregnancy, non-ambulatory status, and an untreated neuropsychiatric or neurological condition. Four individuals withdrew before randomization. The remaining 14 were randomly assigned to a four-week movement intervention — improvisational dance training (n = 7) or a dance-themed movie-watching control (n = 7) — each comprising two 90-minute sessions per week (eight sessions; 12 hours total). The present analyses concern within-session interpersonal coordination rather than the intervention, and therefore draw on every completed hyperscanning session rather than on intervention completion: they are powered by a session-level sample that is larger than, and analytically distinct from, the intervention-completer sample (see Table 1 and Statistical approach). The intervention contrast itself is reported in a companion paper. The protocol was approved by the Virginia Tech Institutional Review Board (#21-798), and all participants provided written informed consent in accordance with the Declaration of Helsinki.

**Table 1.**
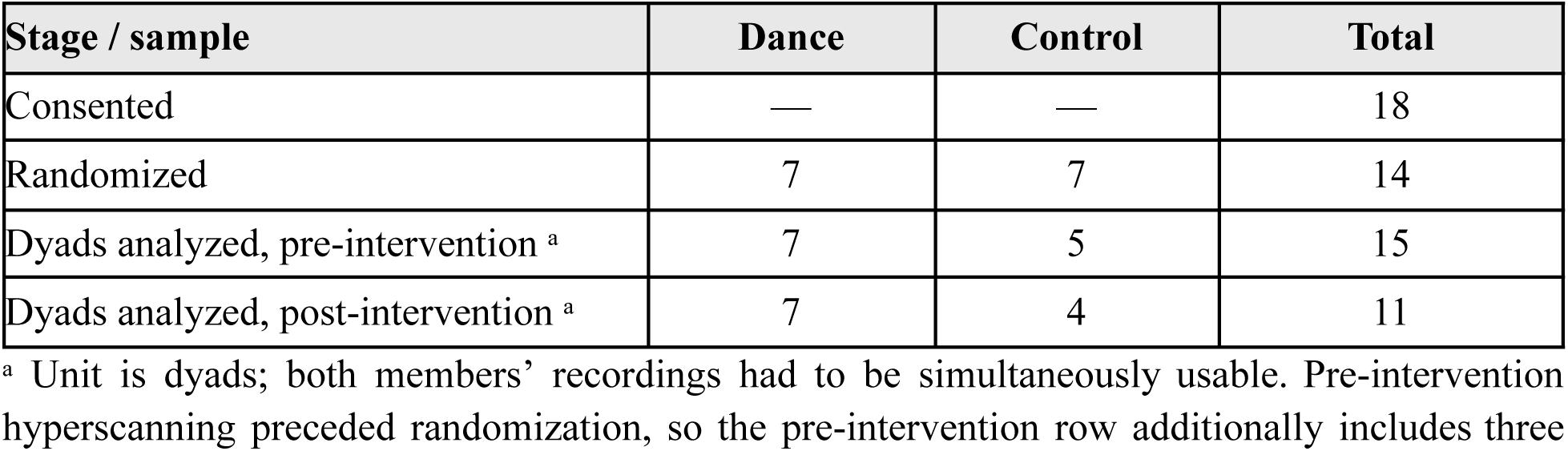
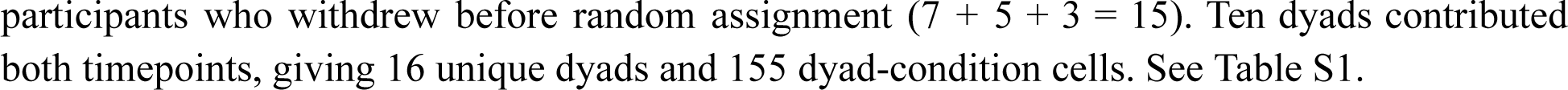
Sample accounting. The within-session coupling analyses reported here are powered by a session-level dyad sample distinct from the intervention-completer sample of the companion paper. Full accounting, including attrition and the intervention-completer rows, is given in Table S1.

| Stage / sample | Dance | Control | Total |
| --- | --- | --- | --- |
| Consented | — | — | 18 |
| Randomized | 7 | 7 | 14 |
| Dyads analyzed, pre-intervention <sup>a</sup> | 7 | 5 | 15 |
| Dyads analyzed, post-intervention <sup>a</sup> | 7 | 4 | 11 |

<sup>a</sup> Unit is dyads; both members' recordings had to be simultaneously usable. Pre-intervention hyperscanning preceded randomization, so the pre-intervention row additionally includes three

Each participant was recorded with a dance instructor using dual-brain hyperscanning EEG across six five-minute conditions that parametrically varied the structure of interpersonal coordination, from none to fully reciprocal: (1) resting baseline (eyes open); (2) direct eye gaze, seated face-to-face at ∼1 m; (3) conversation, planning a shared activity; and three joint-movement conditions — (4) teacher-led mirroring (participant reproduces the instructor’s movements), (5) participant-led mirroring (instructor reproduces the participant’s movements), and (6) movement conversation (the pair initiate, sustain, and end movement in continuous mutual response). This graded design — from imitative synchrony (mirroring) to reciprocal, less-constrained coordination (movement conversation), with non-movement social conditions (eye gaze, conversation) as comparators — is the manipulation on which all analyses rest. Improvisational dance serves here as the elicitation method: it is the most tractable way to evoke sustained, naturalistic, experimentally-gradable dyadic movement, but the questions and measures concern interpersonal movement coordination and its neural signatures in general, not dance per se.

### 2.2 EEG acquisition and preprocessing

Hyperscanning EEG was recorded simultaneously from participant and instructor using two portable 32-channel wet-electrode systems (LiveAmp 32; Brain Products GmbH) at 500 Hz. Recording was continuous across the whole session; preprocessing was performed on the raw continuous data using EEGLAB (MATLAB) with a pipeline optimized for mobile EEG(Böttcher et al., 2026; Rico-Olarte et al., 2025). Data were band-pass filtered (1–45 Hz; zero-phase Hamming-windowed FIR), screened for noisy channels (flat > 5 s, > 4 SD in noise, or inter-channel correlation < 0.8; interpolated), corrected with Artifact Subspace Reconstruction (0.5-s window, 20-SD threshold)(Chang et al., 2020), and re-referenced to the full-rank common average. Independent components were derived by AMICA (single model, 2000 iterations; sample rejection at 3 SD applied five times)(Hsu et al., 2018) and classified with ICLabel(Pion-Tonachini et al., 2019).

Two component-classification thresholds were applied to yield two derived datasets, matched to the demands of different analyses. For channel-level analyses — inter-brain connectivity, channel-level corticokinematic coherence, and the time-resolved co-fluctuation analysis — independent components labeled by ICLabel as > 90% artifact (muscle, eye, heart, line-noise, or other) were removed and the remaining components back-projected to the 32 channels, yielding an artifact-cleaned channel dataset (“IC90”). For component-level analyses that require the source decomposition — specifically the sensorimotor localization of corticokinematic coherence, which is defined over dipole-clustered components — the decomposition retaining components labeled > 50% brain was used (“IC50”), from which the sensorimotor component clusters (precentral, postcentral, and supplementary-motor) were identified. Corticokinematic coherence was therefore computed using a dual-source approach: the channel-level estimate from IC90 (so that it matches the inter-brain analysis) and the component-level localization from the IC50 clusters, each reconstructed from its own dataset’s data. All inter-brain coupling reported here was computed on IC90. Recordings were screened for effective rank as described above; on the IC90 dataset, effective rank ranged from 18 to 31 (median 24), so no recording approached the connectivity floor. Continuous recordings were then segmented into the six conditions using the recorded synchronization markers, preserving component weights.

Two effective-rank criteria were applied, because the two analysis families make different demands. For the inter-brain connectivity analysis, which involves multivariate coupling and is destabilized by rank deficiency, recordings with effective rank (singular values > 10⁻⁶ of the largest) below 15 were excluded(Kim et al., 2023); exclusions were propagated to the dyad, so a pair was dropped for a session if either member failed. Varying the floor confirmed that inclusion did not depend on the threshold: floors of 15 and 18 retained an identical set of recordings, because no recording had an effective rank between 15 and 17, and a floor of 12 additionally retained one participant’s post-intervention recordings (effective rank 14). For corticokinematic coherence and the dyadic movement–neural synchrony analyses, which couple a single kinematic signal to a single neural time series and therefore do not require multivariate rank, a minimal floor of 5 was used to remove only degenerate recordings (one session, effective rank 2). Because retained brain components are the back-projected basis, each recording’s component count equals its effective rank; component indices used below therefore index genuine signal, not null-space.

#### Accelerometry (movement quantification)

Tri-axial accelerometry (dedicated x, y, z channels) was acquired from the same LiveAmp amplifiers as the EEG and segmented with the identical synchronization markers, so movement and neural signals shared a common clock. Accelerometry was extracted before EEG filtering and is therefore unfiltered raw data. For each condition, gravity was removed with a fourth-order Butterworth high-pass filter (0.3 Hz), the three axes were combined into a dynamic-acceleration magnitude (Euclidean norm), and the signal was decimated to 50 Hz(Reis et al., 2014). This magnitude series is the kinematic signal for all movement measures below.

### 2.3 Dyadic movement synchrony

Dyadic movement synchrony between participant and instructor was quantified on the magnitude series with two complementary measures. A windowed cross-correlation (8-s windows, 4-s step, maximum lag ±2 s) yielded, per recording, the mean signed peak correlation (coordination strength) and the median lag at that peak (positive = participant trails instructor). Band-limited coordination was quantified as magnitude-squared coherence averaged over 0.5–3 Hz, the range of coordinated whole-body movement. Because these dance movements are slow, the dominant movement cadence is low (median F0 = 0.53 Hz, IQR 0.44–0.81 Hz).

### 2.4 Partner-specific synchrony

To isolate coupling attributable to the specific pairing rather than to a shared external driver (music, choreography, or simply performing the same task), every synchrony and coupling measure — movement and neural alike — was evaluated against a pseudo-dyad surrogate in which a participant’s signal was paired with mismatched recordings (same timepoint and condition) of other individuals. Partner-specific synchrony was defined as the real value minus the mean of the surrogate distribution. Applying identical surrogate logic to predictor (movement) and other outcomes (neural) ensures that both express pairing-specific coupling, with the shared task structure removed(Burgess, 2013). For the inter-brain analysis, a stricter same-instructor surrogate (participant paired with the same instructor in a different session) was also retained, isolating partner-specificity from stable instructor idiosyncrasy.

### 2.5 Inter-brain coupling (primary analysis)

Inter-brain synchrony was computed with HyPyP (MNE-Python) on the preprocessed 32-channel data. Hardware-synchronized amplifiers shared a common synchronization-pulse train, so the two members’ per-condition recordings were time-aligned; each was epoched into 4-s non-overlapping windows from a shared onset and truncated to common length, with a retained-epoch guard flagging any dyad whose members’ epoch counts diverged by > 10% (catching packet loss that equal-duration cutting would conceal). Dyads whose two members were not recorded in this shared-clock configuration were excluded, because their recordings could not be verifiably time-aligned; this affected one post-intervention dyad, whose session was acquired by Bluetooth streaming rather than to SD card (due to SD card failure). Five connectivity measures were derived from the same complex spectra and averaged over epochs — coherence, phase-locking value (phase only), imaginary coherence (lag-robust phase), amplitude-envelope correlation (amplitude only), and orthogonalized cross-brain AEC (leakage-robust amplitude) — across five bands (theta, alpha, low-beta, high-beta, gamma). For each measure and band, the mean of the full cross-brain block (all participant × instructor channel pairs) indexed inter-brain coupling(Zamm et al., 2024). Partner-specific coupling was the real value minus the mean of the mismatched surrogate. Leakage-robust measures (imaginary coherence, orthogonalized AEC) were treated as primary and zero-lag-sensitive measures as confound checks; because pseudo-dyads reuse individuals and are not mutually independent, permutation p-values were treated as approximate and inference rested on effect sizes and spatial/cross-measure consistency.

### 2.6 Relating movement synchrony to inter-brain coupling

Movement and inter-brain measures were merged by dyad × timepoint × condition, yielding 155 dyad-condition cells from 16 dyads. The primary model was a linear mixed-effects model with partner-specific inter-brain coupling (fit per band) as the outcome. Partner-specific movement synchrony was decomposed by person-mean centering into a between-dyad component (each dyad’s mean, a trait-like tendency to move in synchrony) and a within-dyad component (each cell’s deviation, indexing state-level coordination); both were entered together with condition, timepoint, and instructor as fixed effects and a random intercept for dyad. Because only two instructors delivered sessions, instructor was a fixed two-level covariate (a two-level grouping cannot support a variance component), and the recording-level teacher label was mapped to the human instructor, so this term captured between-instructor variance rather than dyad identity. Predictors and outcome were z-scored (standardized coefficients); significance was assessed using the Benjamini–Hochberg FDR correction across the five bands. As a robustness check isolating the within-dyad effect from any between-dyad confound (instructor, study arm), the same relationship was estimated by OLS with dyad fixed effects and cluster-robust standard errors (by dyad), so the movement coefficient reflects purely within-dyad covariation.

### 2.7 Specificity and artifact-control analyses

Three controls guarded against a movement-artifact account of the inter-brain effect. First, the leakage-robust orth-AEC (which discards zero-lag coupling) and the three phase-based measures were analyzed identically to test whether the association was specific to amplitude-envelope coupling. Second, the analysis was repeated against the stricter same-instructor surrogate. Third, and most directly, movement vigor — each member’s mean dynamic-acceleration magnitude and their interaction (joint movement amplitude) — was added to the model, testing whether the effect reflected the amount of movement (which drives motion/muscle artifact) rather than its temporal coordination.

### 2.8 Corticokinematic Coherence: A Specific Mechanistic Measure

To test whether inter-brain coupling reflects shared tracking of the movement rhythm in the sensorimotor cortex, we computed corticokinematic coherence (CKC) between the accelerometric magnitude and the EEG at the movement fundamental frequency. For each recording the fundamental frequency F0 was estimated as the dominant peak of the acceleration power spectrum within 0.4–3 Hz; magnitude-squared coherence between kinematics and each neural time series was computed by multitaper (8-s segments, 50% overlap, time-bandwidth product NW = 2, three tapers; both signals resampled to a common 100 Hz), and read at F0 and its first harmonic 2·F0 (peak within ±0.15 Hz). These parameters were chosen for the slow dance cadence: an 8-s window yields a 0.25-Hz half-bandwidth, so F0 and 2·F0 remain separable. CKC was computed for four couplings per dyad × condition: within-person (each member’s brain vs. own movement) and cross-person (each member’s brain vs. the partner’s movement). Consistent with the dataset assignment above, channel-level CKC was computed on the artifact-pruned channel dataset and component-level CKC on the dipole-clustered component dataset, each reconstructed from its own decomposition; localization was tested as sensorimotor (precentral/postcentral/SMA cluster components, and a central channel belt: FC1, FC2, C3, Cz, C4, CP1, CP2) versus non-sensorimotor, as disjoint equal-footing sets. Because these analyses couple a single neural signal to movement rather than two multivariate datasets, they used the lenient effective-rank floor (excluding only degenerate recordings) rather than the rank-15 floor applied to inter-brain connectivity; CKC therefore retained all recordings with usable movement and EEG. Statistical baselines were the analytic coherence confidence limit and, for cross-person coupling, the pseudo-dyad surrogate. Within-person coupling additionally used a segment-shuffle null that breaks cross-segment correspondence.

### 2.9 Dyadic movement-neural synchrony at the movement rhythm

As a further specificity test — whether interpersonal coupling takes the form of a shared movement-frequency rhythm linking the two brains — we computed, per dyad × condition at F0, both movement synchrony (participant vs. instructor accelerometer coherence) and neural synchrony (participant vs. instructor brain coherence, averaged over sensorimotor channels and over sensorimotor components), each against the pseudo-dyad surrogate. We then tested, in the same within/between mixed-model framework used for the envelope analysis, whether partner-specific movement synchrony predicted partner-specific neural synchrony. Two robustness analyses accompanied this test: a broadband version summarizing neural coherence across 0.5–4 Hz rather than at F0 alone (guarding against an effect displaced from the exact fundamental), and an exploratory cross-lagged analysis over the ordered conditions within each session (does movement synchrony in one condition predict neural synchrony in the next, and is the reverse direction weaker), fit by dyad-fixed-effects regression and interpreted cautiously given the short six-point series.

### 2.10 Time-resolved body-brain co-fluctuation

To test the association at the timescale at which coordination unfolds, movement synchrony and inter-brain coupling were computed in sliding windows within each condition. In each window, inter-brain coupling was the mean across homologous channel pairs of the correlation between the two members’ amplitude envelopes, summed over the fast bands in which the coupling was observed (high beta, 20–30 Hz; gamma, 30–45 Hz); movement synchrony was the peak of the windowed cross-correlation between the two accelerometry magnitude series, searched over ±2 s. This yielded two time series per dyad-condition, whose within-condition correlation is the co-fluctuation estimate. Confidence intervals on each estimate were obtained by contiguous block bootstrap (45 s blocks, 1,000 resamples) so that temporal autocorrelation among overlapping windows was respected, and inference proceeded from one estimate per dyad-condition rather than treating windows as independent. A partner-specific null was constructed by recomputing the coupling series against mismatched instructor recordings from the same condition and timepoint, giving the surrogate distribution against which real co-fluctuation was compared.

To establish the temporal resolution of the lagged analysis, co-fluctuation and the lagged cross-correlation were computed at three window/step configurations: 15 s windows advancing in 7.5 s steps, 15 s windows advancing in 1 s steps, and 8 s windows advancing in 0.5 s steps. Step size determines the resolution of the lag axis while window length determines how strongly the cross-correlation is smoothed, so the finer configurations locate the peak more precisely without altering the quantity estimated. The peak was located by parabolic interpolation of the three lag values surrounding the maximum, with confidence intervals from 5,000 bootstrap resamples of dyads. All three configurations were run to completion and all are reported.

### 2.11 Statistical approach and software

Inference throughout rested on standardized effect sizes (Hedges *g* for one-sample and contrast tests; standardized β for model slopes) with mixed-effects models (dyad random intercept) as the primary framework and dyad-fixed-effects OLS with cluster-robust standard errors as a within-dyad robustness check. Given the pilot-scale sample (15 pre- and 11 post-intervention dyads) and the non-independence of pseudo-dyad surrogates, permutation *p*-values were treated as approximate, and interpretation emphasized effect size, direction, and convergence across measures and analyses; null results are reported as such rather than as absence of evidence. Consistent with this estimation-focused approach, we did not conduct an a priori power analysis for a fixed effect size; instead, a sensitivity analysis indicated that the primary within-dyad design was adequately powered to detect standardized slopes of β ≥ 0.24 at 80% power (α = 0.05, two-sided), given the observed data structure. The primary effect (β = 0.54, 95% CI [0.37, 0.71]) exceeded this threshold, with its interval excluding both the null and small effects. For the time-resolved co-fluctuation analysis, the pseudo-dyad surrogate provides the null distribution directly, so inference does not rely on a threshold-power assumption. Multiple comparisons were controlled by Benjamini–Hochberg FDR within each analysis family. Analyses were conducted using MATLAB (EEGLAB with DIPFIT and ICLabel) and Python (MNE-Python and HyPyP; statsmodels; SciPy). Analysis code is available at https://github.com/embodiedbrainlab/Dance-on-the-Brain; raw, de-identified EEG or behavioral data are available from the corresponding author upon request.

## 3. RESULTS

### 3.1 Sample and analysis structure

Of 18 adults who consented, four withdrew before randomization, and 14 were randomized to a four-week movement intervention — improvisational dance training (n = 7) or a dance-themed movie-watching control (n = 7). Pre-intervention hyperscanning was conducted at baseline, prior to randomization, so three participants who subsequently withdrew nonetheless contributed a usable pre-intervention session; these are retained because the within-session coupling analyses do not depend on intervention assignment or completion. Because the present analyses concern within-session interpersonal coordination rather than the intervention, they draw on every completed hyperscanning session and are therefore powered by a session-level sample that is analytically distinct from the intervention-completer sample reported in the companion paper (Table 1; full accounting in Table S1). A dyad entered analysis only if both members’ recordings were simultaneously usable after exclusions for effective rank and recording quality, so a single-member exclusion dropped the whole dyad — the reason the analyzed sample here is defined at the dyad rather than the participant level. Sixteen unique participant–instructor dyads contributed at least one usable session: 15 at pre-intervention and 11 at post, of which 10 contributed both. Although modest in dyad count, the design is dense within each dyad: dual-brain EEG spanned the six graded conditions to yield 155 dyad-condition cells (90 pre, 65 post), 775 dyad-minutes of analyzed dual-brain recording, and 6,155 sliding windows for the time-resolved analysis; across five connectivity measures and five frequency bands, this gave 3,875 partner-specific inter-brain coupling estimates

### 3.2 The paradigm created graded partner-specific movement coordination

Before asking whether interpersonal movement synchrony relates to inter-brain coupling, we confirmed that the paradigm produced the intended coordination. Windowed cross-correlation of the two members’ movement-magnitude series recovered the designed lead–lag structure: in teacher-led mirroring (follow) the participant’s movement trailed the instructor’s (median lag +0.12 s), whereas in participant-led mirroring (lead) the lead reversed (-0.10 s); the within-dyad follow-versus-lead difference was robust (mean 0.22 s; paired t = 4.57, p < 0.0001; 24/30 dyad-sessions in the expected direction) (Figure 1A). Partner-specific movement coherence — real minus pseudo-dyad surrogate — was significantly positive and concentrated in the movement conditions (follow, lead, and improvisation, each p ≤ 0.0001 versus surrogate; Figure 1B), confirming that coordination was specific to the actual pairing rather than to shared music.

**Figure 1.**
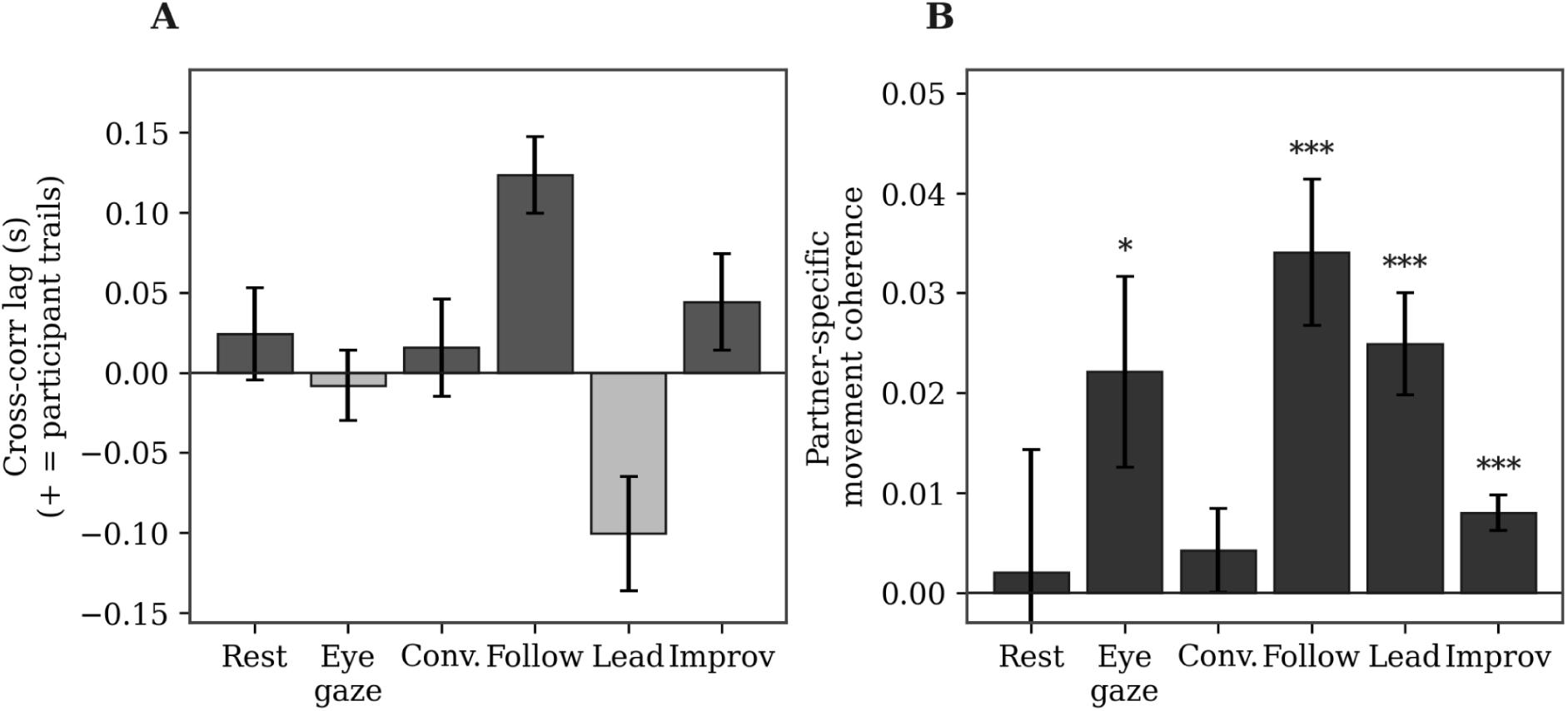
The paradigm elicits graded, partner-specific interpersonal movement coordination. (A) Cross-correlation lead–lag reverses between teacher-led and participant-led mirroring. (B) Partner-specific movement coherence peaks in the movement conditions. Error bars, ±1 SEM across dyad-sessions.

### 3.3 Inter-brain synchrony is partner-specific, amplitude-based, and fast-band

Having established that the paradigm produces interpersonal movement coordination, we next asked whether inter-brain synchrony itself is present above chance pairing — the second phenomenon that the central analysis will relate to movement. Comparing real dyads against same-instructor pseudo-dyads (the strictest surrogate, holding both individuals’ identities constant and removing only real-time simultaneity), partner-specific inter-brain coupling was present, but only in a specific form. Amplitude-based coupling was significantly elevated in the fast bands — envelope correlation reached z = 0.50 in gamma (p = 0.0002) and z = 0.31 in high-beta (p = 0.005), with the leakage-robust orthogonalized measure equally strong (gamma z = 0.54, p < 0.0001; high-beta z = 0.35, p = 0.002), indicating the coupling is not an artifact of volume conduction. In contrast, none of the three phase-based measures (coherence, PLV, imaginary coherence) exceeded surrogate in any band, and amplitude coupling in the slow bands (theta, alpha, low-beta) was null (Figure 2; full measure × band values in Table S2; condition-resolved pattern is shown in Figure S1). Restricting the cross-brain average to homologous channel pairs yielded the same pattern — significant amplitude coupling in high-beta (envelope correlation z = 0.31, orth-AEC z = 0.31) and gamma (z = 0.56 and 0.58), with all phase measures null in every band — indicating that the effect does not depend on whether coupling is averaged over the full cross-brain block or over matched pairs alone (Table S3). Inter-brain synchrony in these dyads is therefore specifically a fast-band amplitude phenomenon — present as co-modulation of high-beta and gamma envelopes between the two brains, and absent as phase synchrony. This independently motivates the focus of the following analysis on amplitude coupling and fast bands: the coupling that exists is of that kind.

**Figure 2.**
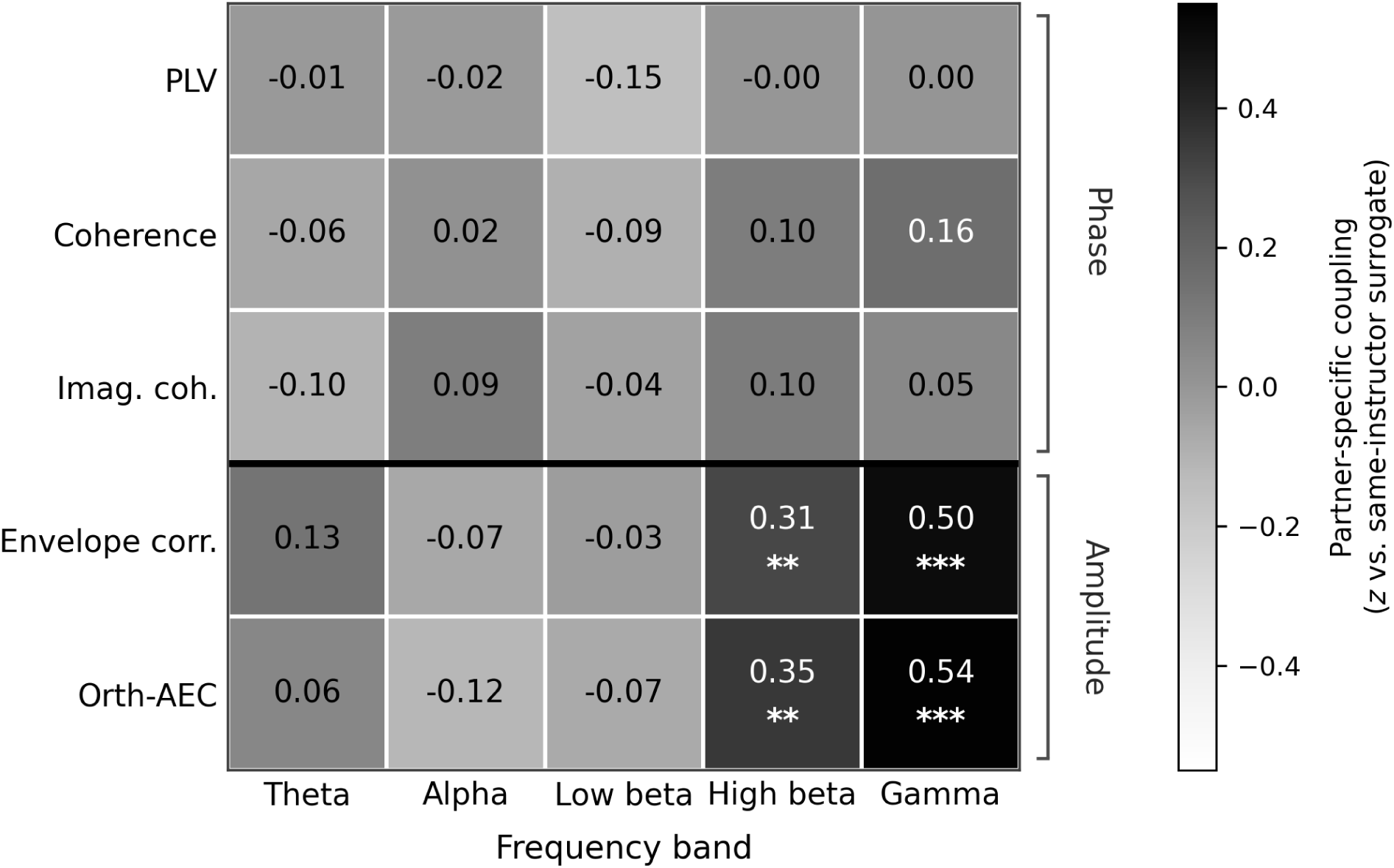
Inter-brain synchrony is partner-specific, amplitude-based, and fast-band. Partner-specific coupling by connectivity measure and frequency band, expressed as the mean z of real dyads against a pooled same-instructor pseudo-dyad null across 155 dyad-condition cells from 16 dyads. Amplitude measures (below the divider) show coupling in high-beta and gamma; phase measures (above the divider) show no coupling in any band, and neither measure class couples in the slow bands. Asterisks denote one-sample t-tests against zero: **p < .01, ***p < .001; Benjamini–Hochberg FDR-corrected values are given in Table S2.

The coupling showed no regional specificity. Partner-specific gamma envelope coupling was significant in 35 of 36 participant-region by instructor-region pairs and did not differ among them (one-way across pairs, F = 0.69, p = 0.914; range of z across pairs = +0.21 to +0.56). Homologous pairs, in which a participant’s region was compared with the instructor’s corresponding region, were no stronger than heterologous pairs (z = +0.39 vs. +0.34, p = 0.317). The same uniformity held for high-beta envelope correlation (F = 0.84, p = 0.731) and for the leakage-corrected orthogonalized measure in gamma (F = 0.83, p = 0.742). Coupling was therefore spatially distributed rather than confined to particular region pairs (Figure S2).

### 3.4 Interpersonal movement synchrony predicts inter-brain amplitude coupling

The primary analysis asked whether partner-specific movement synchrony predicts partner-specific inter-brain coupling, within dyads. Using the cross-correlation peak as the movement-synchrony measure — the time-domain index best suited to the slow, non-periodic coordination these dyads produced — within-dyad movement synchrony strongly predicted inter-brain amplitude-envelope coupling in the gamma band (standardized β = 0.54, 95% CI [0.37, 0.71], p < 0.0001, FDR q < 0.001; envelope correlation). The effect increased monotonically across frequency — theta (β = 0.16, n.s.) through alpha (0.21), low-beta (0.36), high-beta (0.44) to gamma (0.54) — and held for the leakage-robust orthogonalized measure (orth-AEC, gamma β = 0.49, q < 0.001), indicating the coupling is not an artifact of volume conduction (Figure 3A; Table 2; within-dyad slopes for all measures and bands in Table S4). Critically, the association was undiminished by movement amount: adding each member’s movement vigor and their interaction left the effect unchanged (β = 0.54 → 0.55; all vigor terms n.s.), and it survived a dyad fixed-effects estimator with cluster-robust standard errors (β = 0.54, p < 0.0001), isolating purely within-dyad, state-level covariation (Figure 3B).

**Figure 3.**
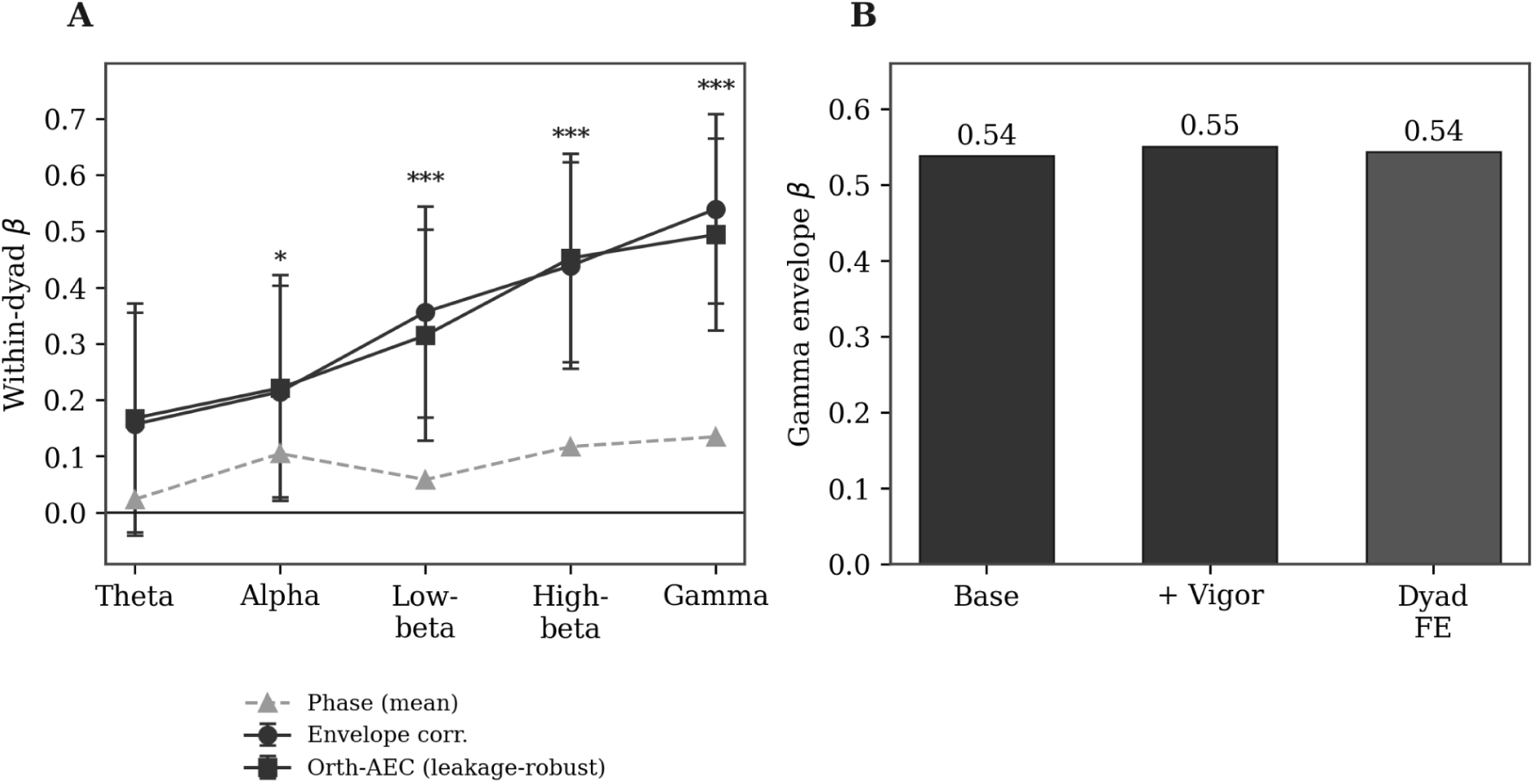
Interpersonal movement synchrony predicts inter-brain amplitude coupling, specifically in fast bands. (A) Within-dyad standardized β by frequency band for amplitude measures (envelope correlation, orth-AEC) versus the mean of phase measures; error bars, 95% CI. (B) The gamma envelope effect is unchanged by movement-amount control and by dyad fixed effects.

**Table 2.**
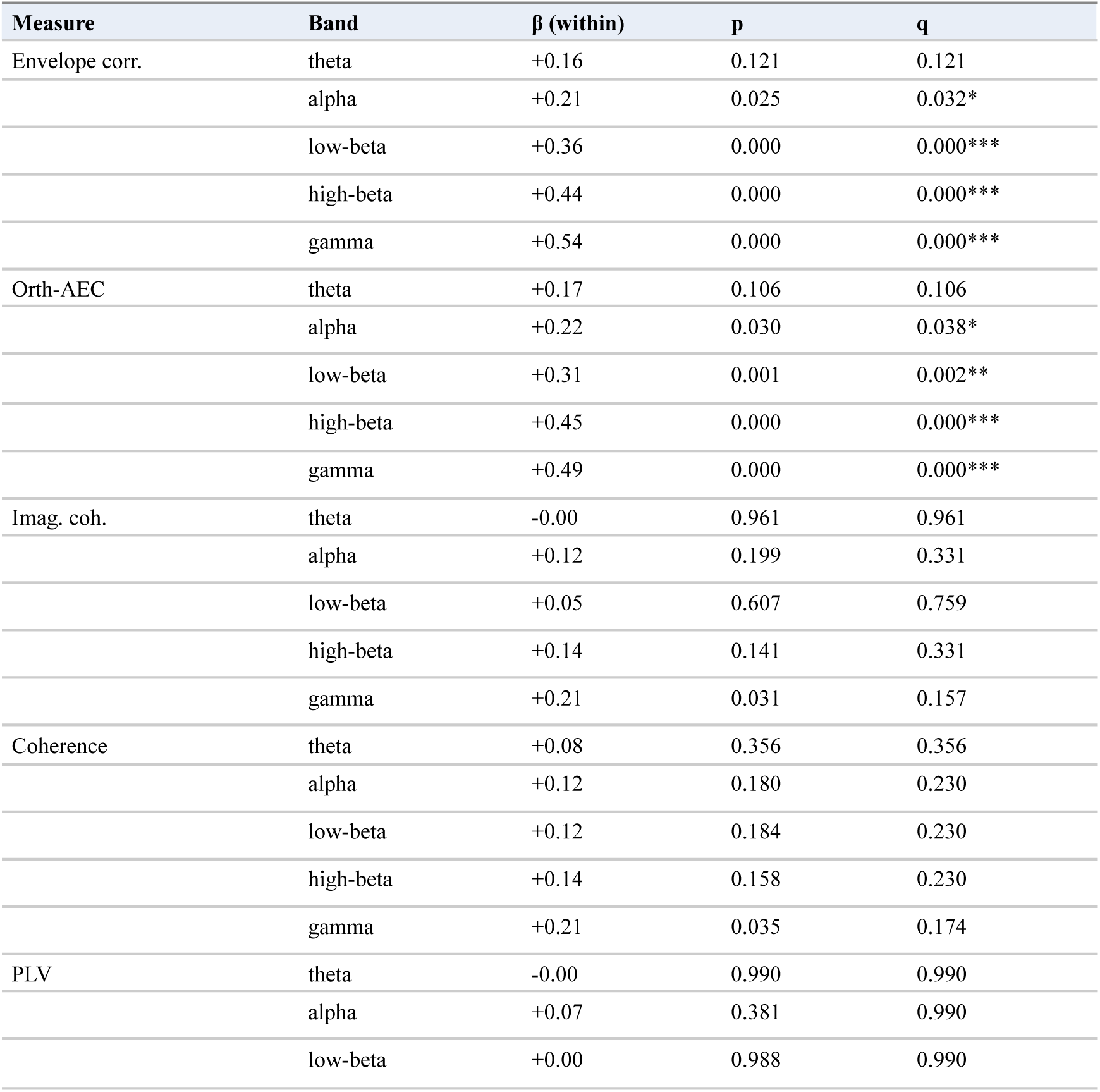
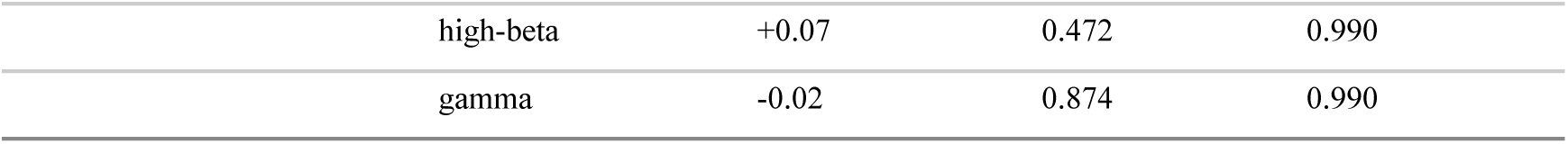
Within-dyad movement-synchrony → inter-brain coupling by measure and band (cross-correlation-peak predictor). Standardized β; q, Benjamini–Hochberg FDR across bands within measure.

The pre-specified secondary measure — band-limited movement coherence averaged over 0.5–3 Hz — showed the same direction and specificity but a smaller effect (gamma envelope β = 0.19, p = 0.016). A post hoc analysis, prompted by the observation that the dominant movement cadence was slower than anticipated (median F0 = 0.53 Hz, IQR 0.44–0.81 Hz), indicated that this attenuation is at least partly an artifact of the fixed band: in 40% of recordings the movement fundamental fell below the 0.5 Hz floor and was therefore excluded from the coherence estimate. Recomputing movement coherence at each recording’s own fundamental — the same per-recording F0 used in the corticokinematic analyses — recovered an intermediate effect (gamma envelope β = 0.26, p = 0.0006, dyad fixed effects). The three operationalizations thus order by how closely each tracks the movement actually present: strongest for the cross-correlation peak, which makes no spectral assumption (β = 0.54); intermediate for coherence read at the true fundamental (β = 0.26); weakest for coherence over a fixed band that omits that fundamental in a substantial minority of recordings (β = 0.19). All three agree in sign, spectral profile, and amplitude-versus-phase specificity. We therefore report the effect as robust in direction and specificity across operationalizations, and strongest with the time-domain measure (Table S4)..

### 3.5 The coupling is not sensorimotor rhythm tracking (control)

Two specificity analyses tested whether the effect could instead reflect shared tracking of the movement rhythm. First, corticokinematic coherence (CKC) between movement and each brain, evaluated at the movement fundamental, confirmed that individual cortex does track the movement rhythm above chance pairing in every condition (within-person Hedges g = 0.43–1.10; Figure 4A) — but with two features that exclude a sensorimotor-tracking account of the inter-brain effect. The coupling did not localize to sensorimotor cortex: sensorimotor and non-sensorimotor components did not differ (IC-level g = 0.02, p = 0.81; Figure 4B; condition-wise CKC, localization, and cross-person controls in Table S5). Additionally, it was not selective to the movement conditions — CKC was as high or higher during the near-still social conditions (eye gaze, conversation) as during mirroring and improvisation, unlike the inter-brain effect, which is organized by movement coordination specifically. Cross-person CKC — one member’s cortex versus the partner’s movement — was not above surrogate (g = 0.11, p = 0.13). Second, dyadic neural synchrony at the movement frequency was essentially absent: although movement synchrony was strongly partner-specific (g = 0.40, p < 10⁻⁶), partner-specific brain–brain coherence at that frequency was negligible over sensorimotor channels (g = 0.19) and null over sensorimotor components (g = 0.04, p = 0.65), as movement synchrony did not predict it (within-dyad β = 0.005, p = 0.96; Table S5). Together, these establish that the inter-brain coupling organized by movement synchrony is fast-band amplitude co-modulation, not a shared low-frequency rhythm and not proprioceptive sensorimotor tracking.

**Figure 4.**
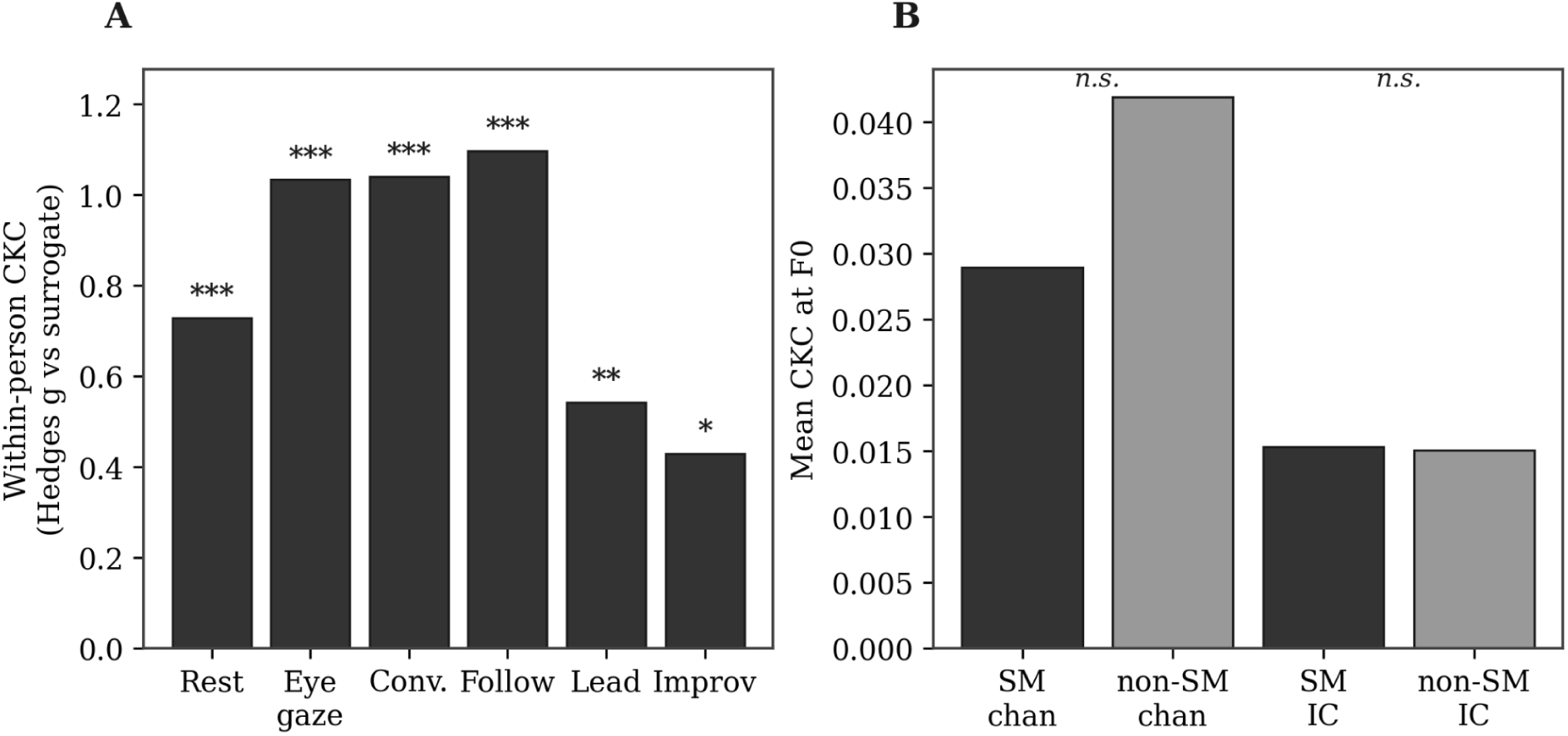
Corticokinematic coherence is a control, not the mechanism. (A) Within-person CKC exceeds surrogate in all conditions but is not selective to the movement conditions as it is high during near-still social conditions too. (B) CKC does not localize to sensorimotor cortex (sensorimotor vs non-sensorimotor, n.s.).

### 3.6 Body-brain co-fluctuations with interactions

The condition-level analysis relates one movement-synchrony value to one coupling value per five-minute condition. To test the hypothesis at the timescale it concerns — whether, moment-to-moment within an interaction, tighter interpersonal movement coordination accompanies tighter inter-brain coupling — we computed both quantities in sliding 15-second windows and measured their within-condition co-fluctuation, with partner-specificity established against a pseudo-dyad surrogate and temporal autocorrelation handled by block bootstrap. Movement synchrony and inter-brain fast-band amplitude coupling co-fluctuated within interactions well above the pseudo-dyad surrogate: mean within-condition r = 0.11 for real dyads versus 0.007 for surrogates (which did not differ from zero, p = 0.13), giving a partner-specific co-fluctuation of 0.099 (t = 5.92, p < 0.0001; real exceeded surrogate in 102 of 155 dyad-conditions; Figure 5A). Because the surrogate — the same participant’s brain against mismatched partners’ movements — showed no co-fluctuation, the effect cannot be attributed to shared task structure or to the windowing itself. Whether either channel led the other could not be resolved. At the original 7.5 s window step the lagged analysis gave no advantage to movement leading neural coupling (r = 0.022) over the reverse (r = 0.025; Figure 5B, 5D), but the resolution of that analysis is far coarser than the timescale at issue: the lead–lag structure of interpersonal movement coordination in these same dyads was 0.12 s. Recomputing the lag profile at finer resolution narrowed the estimate substantially. Across window/step configurations of 15 s/7.5 s, 15 s/1 s, and 8 s/0.5 s, the sub-grid peak moved from -0.07 s to +0.22 s to +0.17 s while the dyad-level bootstrap 95% confidence interval narrowed from 1.38 s to 0.91 s to 0.51 s (Figure S3). The finest configuration placed the peak at +0.17 s in the direction of movement leading neural coupling, with 95% CI [-0.04, +0.47]. Conservative per-dyad tests did not support a lead, however: profile asymmetry across ±0.5–3 s was non-significant (11 of 16 dyads positive, p = 0.55), and per-dyad peak locations were evenly split (8 of 16, p = 1.00). These analyses therefore exclude lags greater than approximately 0.5 s in either direction, and are consistent with — but do not establish — movement synchrony preceding inter-brain coupling.

**Figure 5.**
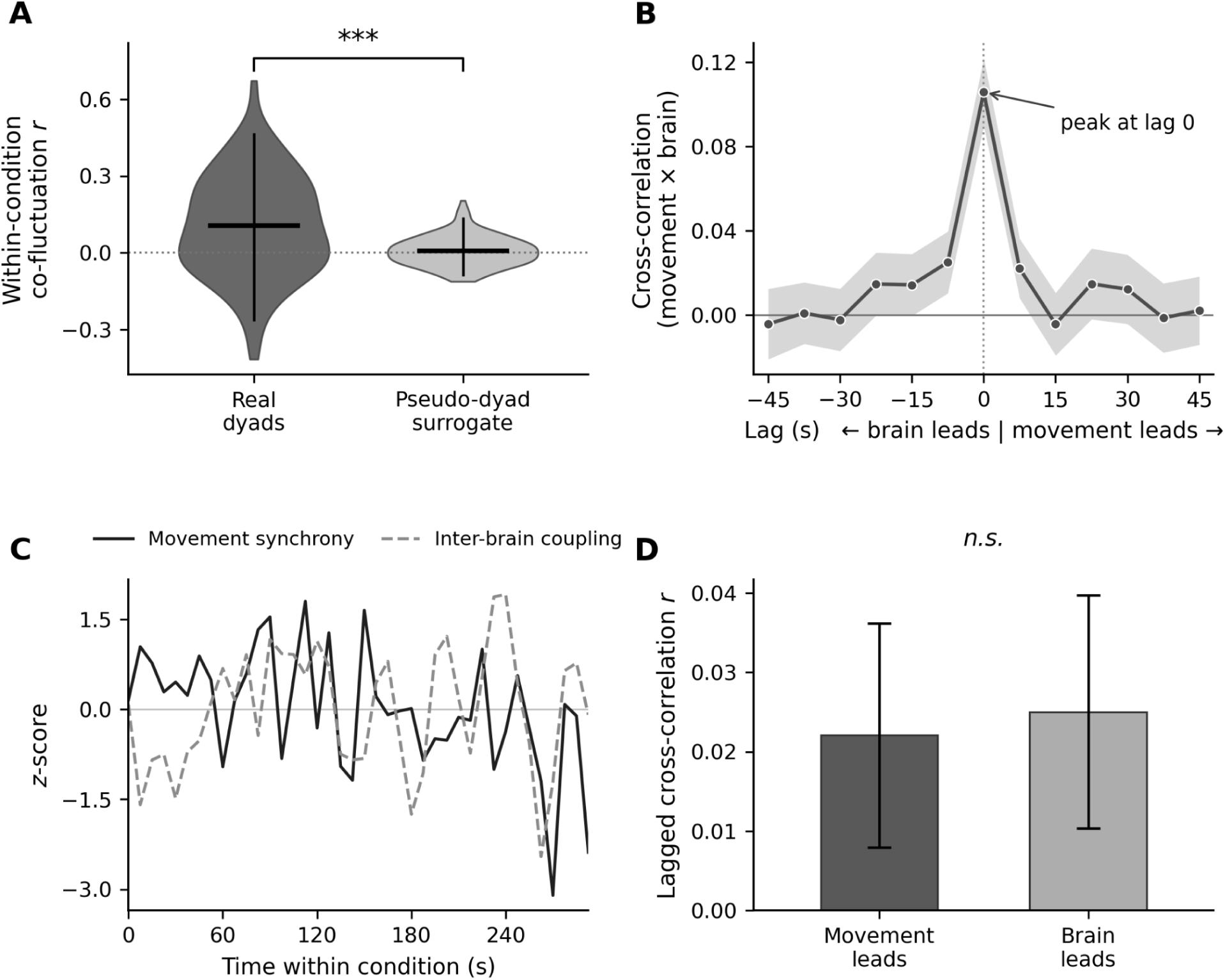
Interpersonal movement synchrony and inter-brain coupling co-fluctuate within interactions, partner-specifically and simultaneously. (A) Within-condition co-fluctuation (movement synchrony × inter-brain fast-band coupling, sliding 15-s windows) for real dyads versus pseudo-dyad surrogates; partner-specific co-fluctuation = 0.099, p < 0.0001. (B) Cross-correlation of the two window series averaged across all dyad-conditions, peaking sharply at lag 0 (simultaneous co-fluctuation). (C) Representative dyad: z-scored movement-synchrony and inter-brain-coupling series over time within one condition. (D) Lagged analysis shows temporal symmetry — no evidence that movement leads neural coupling or the reverse.

## 4.0 DISCUSSION

Across a graded series of interpersonal conditions, we found that interpersonal movement coordination organizes inter-brain coupling. Two phenomena were each present and specific to the interacting pair: movement synchrony, which varied systematically with the structure of the task, and inter-brain synchrony, which took the specific form of amplitude-envelope co-modulation in the high-beta and gamma bands and was absent from phase-based measures. Within dyads, the degree of movement coordination predicted the strength of this fast-band amplitude coupling, an association undiminished by the amount of movement and by dyad-level differences. The two analyses converged on the same spectral signature: the surrogate contrast and the movement-synchrony prediction both located the effect in the fast amplitude envelopes and both strengthened monotonically toward gamma. Time-resolved analysis showed that the two quantities co-fluctuated moment-to-moment within interactions, rising and falling together and partner-specifically, at a lag smaller than half a second but without a resolvable direction of influence. Two specificity controls, corticokinematic coherence and dyadic synchrony at the movement frequency, excluded the most immediate alternative explanations. Together these results identify what organizes inter-brain coupling here: not the mere presence of a partner, and not how much two people move, but how precisely they move together.

### 4.1 Movement coordination organizes inter-brain coupling

The central result is that inter-brain coupling is not a fixed property of two people being together but is graded by the degree to which their movements are coordinated. This speaks directly to a longstanding interpretive problem in the hyperscanning literature: when two brains appear synchronized, the coupling may reflect genuine interaction or may be produced by shared sensory input, common task structure, or correlated movement(Schilbach and Redcay, 2025). Our design addresses this in three converging ways. Coupling was evaluated against pseudo-dyad surrogates that hold the task constant while breaking the real-time pairing, so that what remains is specific to the actual partners rather than to shared structure. It was shown to scale with a directly measured behavioral quantity, interpersonal movement synchrony, rather than merely to co-occur with interaction. It also survived the inclusion of movement vigor, dissociating coupling organized by the temporal coordination of movement from coupling that merely accompanies the amount of movement (and its attendant artifact). The within-dyad, state-level nature of the association is particularly informative: it holds after removing each dyad’s average tendency, reflecting moment-to-moment changes in coordination within a pair, not stable differences between pairs.

### 4.2 An amplitude phenomenon, not phase synchrony

That the coupling appeared in amplitude-envelope measures and not in any phase-based measure has implications for the nature of inter-brain coupling. Much hyperscanning work operationalizes inter-brain synchrony as phase alignment(Czeszumski et al., 2020; Zimmermann et al., 2024), yet phase measures here (coherence, the phase-locking value, and imaginary coherence) showed no partner-specific coupling in any band, whereas envelope correlation and its leakage-robust orthogonalized variant showed reliable coupling in high-beta and gamma. This converges with recent evidence placing spontaneous inter-brain coupling in the beta and gamma envelopes(Koul et al., 2023). The convergence is itself informative: that study selected an amplitude measure in advance, on the grounds that phase estimates are unreliable when band-limited power is low(Burgess, 2013; Hipp et al., 2012). The present analysis supplies the comparison that this choice anticipates — five measures estimated from the same complex spectra, with coupling appearing only in the amplitude-based two. What has been a principled analytic assumption is here an empirical dissociation.

Amplitude-envelope coupling reflects co-modulation of the strength of band-limited activity between the two brains rather than alignment of its instantaneous phase(Engel et al., 2013; Hipp et al., 2012), and is thought to index coordinated fluctuations in cortical excitability and engagement(Engel et al., 2013). The band specificity reinforces this reading: high-beta and gamma amplitude track moment-to-moment attentional and arousal-related engagement, the very processes that reciprocal movement coordination demands. This pattern of amplitude synchronization and phase indifference has also been observed to be modulated by interpersonal aspects of a task. One study found that synchronous inter-brain amplitude was positively correlated during collaborative game tasks, although this was only significantly observed in theta and delta bands(Liu et al., 2021). Due to the highly cooperative nature of improvisational dance, our findings support these results and suggest that amplitude-based coupling is strongly related to coordinated movement. Gamma amplitude-envelope coupling has also been observed in dueting pianists, during still breaks in performance, providing further support that this coupling is the result of inter-brain synchrony rather than other muscle artifacts(Gugnowska et al., 2022). The convergence of two independent analyses, the surrogate contrast and the movement-synchrony predictor, on the same high-frequency amplitude signature makes it unlikely that this pattern is an incidental feature of one analytic choice. It also suggests that studies restricting inter-brain analyses to low-frequency phase coupling may miss coupling that is present in the amplitude of faster rhythms.

The spatial uniformity of the coupling constrains what can be generating it. Interaction between specific cortical systems would be expected to concentrate in particular region pairs, and homologous pairs in particular; neither was observed. A more parsimonious reading is co-modulation of a global fast-band amplitude state within each brain — the kind produced by shifts in arousal, effort or engagement — becoming correlated between partners as they coordinate. This fits the other features of the effect: global excitability shifts modulate the amplitude envelope rather than aligning instantaneous phase, which is what the amplitude-versus-phase dissociation shows, and broadband gamma amplitude is among the most reliable indices of cortical engagement, which is consistent with the monotonic strengthening toward higher frequencies. It should be noted that 32-channel scalp EEG reduced to six regional aggregates has limited spatial resolution, so uniformity at this scale does not exclude localized underlying sources; what it does exclude is the strong form of the claim that coupling reflects interaction between specific, identifiable cortical systems.

### 4.3 Real-time coupling without a leader

The time-resolved analysis extends the condition-level result to the timescale at which coordination actually unfolds. Within a single interaction, windows of tighter movement coordination coincided with windows of stronger inter-brain amplitude coupling, above what mismatched partners produced. This is a stronger form of the central claim than the condition-level association: it shows that the two quantities track one another dynamically, not merely that dyads who coordinate more also couple more on average.

The aggregate cross-correlation peaked near zero lag, but this should be read as a bound on the timescale of any lead rather than as evidence of true simultaneity. Progressively finer analyses narrowed the interval around the peak by nearly a factor of three, converging on an estimate of approximately +0.17 s in the direction of movement leading neural coupling, while conservative per-dyad tests remained null. What these data establish is that neither channel leads the other by more than about half a second; a lead at the 0.1–0.2 s scale characteristic of interpersonal movement coordination is neither demonstrated nor excluded. This matters for comparison with prior work: synchronized smiling has been reported to precede gamma envelope coupling by approximately 200 ms, with behavioral synchrony Granger-causing inter-brain coupling more strongly than the reverse(Koul et al., 2023). A lead of that magnitude would fall inside our confidence interval and would not have been detectable here, so the apparent discrepancy between the two findings may be one of resolution rather than of phenomenon. It is also possible that sustained reciprocal movement genuinely lacks the event structure that permits a lead: unlike a smile, which has an onset that a neural response can follow, continuous mutual adjustment offers no discrete moment for one channel to precede the other. What the present analysis establishes is that body coupling and brain coupling rise and fall together within interactions, partner-specifically and at a lag smaller than half a second.

### 4.4 Ruling out sensorimotor rhythm tracking

Several analyses exclude the most obvious alternative account: that both brains are independently entrained to a shared movement rhythm. Corticokinematic coherence confirmed that each individual’s cortex tracked the rhythm of its own movement, but this tracking did not localize to sensorimotor cortex and was not selective to the movement conditions — it was as strong during near-still social conditions as during joint movement, unlike the inter-brain effect, which was organized by movement coordination specifically. The inter-brain coupling therefore does not reduce to two brains independently tracking a shared movement rhythm through sensorimotor entrainment. Cross-person coherence was weak and asymmetric: the instructor’s movement rhythm was tracked by the participant’s cortex above surrogate (mean difference = 0.0026, p = 0.013), consistent with the participant attending to and following the instructor, whereas the reverse direction was null (p = 0.149). This tracking cannot account for the inter-brain effect. It occurs at the movement fundamental — a median of 0.53 Hz — whereas the inter-brain coupling is confined to high-beta and gamma amplitude envelopes, and dyadic neural synchrony at the movement frequency was itself negligible and unpredicted by movement synchrony. One partner’s cortex tracking the other’s movement rhythm does not, in these data, produce coupled neural activity at that rhythm. What remains after these controls is coupling in the amplitude of fast cortical rhythms, organized by the temporal coordination of movement and specific to the interacting pair — a signature that the most immediate artifactual and rhythm-tracking accounts do not explain.

### 4.5 Relationship to prior work

These findings connect to a body of hyperscanning work in which inter-brain coupling is partner-specific and shaped by nonverbal exchange. Inter-brain synchrony has been reported to depend on the specific pairing rather than on shared task alone — for example, temporoparietal coupling emerging among romantic couples but not strangers, and tracking the exchange of social gaze rather than verbal content(Kinreich et al., 2017).

The closest precedent to the present work is Koul et al. (2023), who recorded dual EEG alongside automated behavioral tracking in dyads instructed only to look at one another, and showed that spontaneous body movement, smiling, and eye contact each synchronized between partners and predicted inter-brain coupling(Koul et al., 2023). The present results extend that finding in four respects. First, coordination was manipulated rather than observed: six conditions graded interpersonal coordination from stillness to fully reciprocal joint movement, so the association reflects coupling tracking a designed gradient rather than covarying with incidental behavior. Second, the design imposes and reverses roles, and the lead–lag structure of the movement measure flipped sign according to which partner was instructed to lead, confirming that it registers one partner responding to the other rather than both responding independently to the shared music. Third, movement here was sustained and whole-body rather than incidental to a seated task, placing the coupling in precisely the regime where movement-artifact accounts are most pressing. Fourth, synchronized accelerometry allowed movement amount to be entered into the model alongside movement coordination, dissociating coupling organized by the temporal structure of movement from coupling that merely accompanies its magnitude — a control that topographic and preprocessing-based arguments approximate but cannot replace.

Our results add a behavioral organizing variable to this picture: not only is coupling partner-specific, its strength is graded by how tightly the partners coordinate their movements, moment to moment. They also complement the small hyperscanning literature on coordinated action, in which inter-brain coupling rises with the demands of maintaining coordination(Lender et al., 2023) and tracks coordination-relevant features of partnered movement(Bigand et al., 2025). Interpreted through a second-person framework, in which inter-brain synchrony is a relational process that emerges during real-time interaction rather than a similarity between isolated brains(Schilbach and Redcay, 2025), the present results identify interpersonal movement coordination as one concrete process through which that relational coupling is expressed.

### 4.6 Limitations

Several limitations qualify these conclusions. The sample is modest in the number of dyads, and although the design is dense within each pair, inference is at the dyad level; the effect-size estimates, while precise enough to exclude small effects, warrant confirmatory replication at larger scale. The core relationships are correlational: movement coordination and inter-brain coupling covary, but the temporally bounded co-fluctuation provides no evidence of a causal direction, and establishing one will require designs that perturb movement coordination or neural activity directly. Scalp EEG limits spatial inference. Although coupling was uniform across all region pairs tested, 32 channels aggregated to six regions cannot resolve whether that uniformity reflects a genuinely global process or the spatial smearing of localized sources; higher-density recording, or modalities with better source separation, would be required to distinguish these. Because conditions were presented in a fixed order of increasing coordination, we cannot fully separate condition from time-on-task, a caveat that applies with particular force to any measure sensitive to slow drift. Additionally, pseudo-dyad surrogates, while the appropriate control for partner-specificity, are not fully independent of the real data, so the associated p-values are approximate and were interpreted alongside effect sizes and convergence across analyses rather than as thresholds. Finally, because each participant was recorded with one of two dance instructors, the same instructors recur across dyads; we modeled instructor as a fixed covariate and defined partner-specificity against a same-instructor surrogate, so the reported coupling reflects the specific participant–instructor pairing rather than stable instructor idiosyncrasy, but the limited number of instructors means instructor-related variance is estimated coarsely and generalization to participant–participant dyads remains to be tested.

A further limitation concerns the physiological origin of the fast-band coupling. High-beta and gamma EEG are vulnerable to electromyographic (EMG) activity from facial, neck, and scalp muscles, which accelerometry does not fully capture(Gorjan et al., 2022). Our design mitigates the most direct movement-artifact account in several ways: the leakage-robust orthogonalized measure removes zero-lag shared components; the effect was undiminished by each member’s movement vigor and their interaction, dissociating it from the amount of gross body movement that drives motion and muscle artifact; and the coupling appeared specifically in the amplitude of fast rhythms rather than as a shared movement-frequency rhythm, which a crude artifact account would not predict. These controls exclude coupling that scales with the magnitude of whole-body movement, but they cannot fully exclude fine-motor or facial EMG that is itself coordinated between partners — for example, correlated facial expression, subtle head stabilization, or breathing that co-varies with the coordination of interaction rather than with its gross amount. Independent component analysis with ICLabel-based rejection removed components confidently identified as muscle, but this does not settle the question: the conservative rejection threshold retains sub-threshold muscle activity by design, a 32-channel montage cannot resolve the many quasi-independent generators of facial and neck EMG, and ICA separates sources only insofar as they are statistically independent — an assumption violated precisely when peripheral and cortical activity co-vary with the same interpersonal dynamics. Importantly, such coordinated peripheral activity would not be a trivial measurement artifact but a genuine, neurally driven dyadic phenomenon; the open question is whether the coupling we observe is cortical, peripheral (neuromuscular), or an integrated cortico-muscular process spanning the two bodies. Distinguishing these will require converging evidence: high-density montages capable of resolving muscle generators, simultaneous face and neck EMG entered as nuisance regressors, and recording modalities less sensitive to muscle (e.g., source-resolved MEG or intracranial data). We therefore describe the effect as interaction-specific, movement-organized fast-band coupling of neural or neuromuscular origin, and treat the integrated-system interpretation as a hypothesis these data motivate rather than establish.

### 4.7 Conclusion

When two people coordinate their movements, their brains couple — specifically through co-modulation of high-beta and gamma amplitude, in a manner graded by the tightness of their coordination, partner-specific, and expressed moment-to-moment in real time. By measuring movement and brain activity together while testing the coupling against the alternatives of shared stimulation and motion artifact, the present study identifies interpersonal movement coordination as a concrete behavioral process that organizes inter-brain coupling, and specifies that coupling as a fast-band amplitude phenomenon. Interpersonal coordination, long recognized as a foundation of human social life, leaves a definable signature in the coupling between brains.

## Acknowledgments

The authors thank Professor Rachel Rugh for guiding the movement exercises and for generously offering her brain for study.

## Funding

This work was supported by the Integrated Translational Health Research Institute of Virginia (iTHRIV) Scholars Program, funded by the National Center for Advancing Translational Sciences of the National Institutes of Health [award numbers UL1TR003015, KL2TR003016]; and the Virginia Tech Institute for Creativity, Arts, and Technology. The funders had no role in study design, data collection and analysis, the decision to publish, or preparation of the manuscript.

## CRedit authorship contribution statement

Noor Tasnim: Data curation, Software, Methodology, Formal analysis, Writing – review and editing. Rachel M. DeLauder: Investigation, Formal analysis, Writing – review and editing. Mackenzie Aychman: Investigation, Data curation, Writing – review and editing. Mary Gahagan: Investigation, Writing – review and editing. Jessica Purevtugs: Investigation, Writing – review and editing. Jesse Newpol: Investigation, Writing – review and editing. Ryan Frank: Investigation, Writing – review and editing. Grace Grizzell: Investigation, Writing – review and editing. Grace Nobriga: Investigation, Writing – review and editing. Sarah Rose: Investigation, Writing – review and editing. Julia C. Basso: Conceptualization, Methodology, Formal analysis, Investigation, Resources, Supervision, Project administration, Funding acquisition, Visualization, Writing – original draft, Writing – review and editing.

## Declaration of competing interests

The authors declare that they have no known competing financial interests or personal relationships that could have appeared to influence the work reported in this paper.

## Ethics statement

The protocol was approved by the Virginia Tech Institutional Review Board (approval #21-798, approved [date]), and all participants provided written informed consent in accordance with the Declaration of Helsinki.

## Data availability

Analysis code supporting the findings of this study is available at https://github.com/embodiedbrainlab/Dance-on-the-Brain. Raw electroencephalographic recordings are not publicly deposited because participants did not consent to public release of identifiable neural and video data; de-identified recordings and derived data are available from the corresponding author on reasonable request, subject to a data use agreement.

## Declaration of generative AI and AI-assisted technologies in the manuscript preparation process

During the preparation of this work the authors used Claude in order to identify proper statistical analysis tools and create analysis code; Undermind in order to conduct a literature review; and Figure Labs to help generate the graphical abstract. After using this tool, the authors reviewed and edited the content as needed and take full responsibility for the content of the published article.

## Notes

https://github.com/embodiedbrainlab/Dance-on-the-Brain

### Competing Interest Statement

The authors have declared no competing interest.

